# Hydraulic retention time drives changes in energy production and the anodic microbiome of a microbial fuel cell (MFC)

**DOI:** 10.1101/2023.05.21.541624

**Authors:** Antonio Castellano-Hinojosa, Manuel J. Gallardo-Altamirano, Clementina Pozo, Alejandro González-Martínez, Jesús González-López

## Abstract

The fish-canning industry generates large quantities of wastewater that typically contains high concentrations of organic matter and salts. However, little is known about the potential valorization of this type of industrial wastewater using the microbial fuel cell (MFC) technology operated in a continuous flow mode. This study investigated the impacts of three different hydraulic retention times (HRT) on the performance, energy production, and prokaryotic and eukaryotic anodic microbiome of an MFC inoculated with activated sludge from a seafood industry and fed with synthetic wastewater that mimics fish-canning effluents. Three consecutive HRTs were studied: 1 day (HRT1), 3 days (HRT3), and 6 days (HRT6) for 30 days, 21 days, and 21 days, respectively. Voltage, current density, and power density were significantly greater at HRT1 compared to HRT3 and HRT6, whereas no differences in coulombic efficiency (CE) were detected among HRTs. Decreases in the efficiency of removal of organic compounds and increases in the abundance of archaeal communities with increased HRT was related to limited energy production at greater HRT. The increased energy production at HRT1 was tightly linked to increased and decreased absolute abundances of bacterial and archaeal communities, respectively. Variations in the HRT significantly impacted the diversity and composition of the prokaryotic community with critical impacts on energy production. The proliferation of known and diverse electroactive microorganisms, such as those belonging to the genera *Geobacter*, *Shewanella*, *Arcobacter*, and *Clostridium*, was related to increased energy production at HRT1. However, HRT3 and HRT6 enhanced the growth of archaeal methanogens (mainly *Methanosarcina* sp.), which negatively impacted current production. The eukaryotic community showed less sensitivity to changes in HRT and no significant impact on current production. The carbon oxygen demand and organic matter removal % increased from approximately 20% at HRT1 to almost 60% at HRT6. This study shows there is a critical balance between the HRT and prokaryotic microorganisms contributing to organic removal rate and increases and decreases in energy production in an MFC treating wastewater from the fish-canning industry and operated in a continuous mode.

## 1. Introduction

The processing of seafood produces a large quantity of wastewater through various operations such as washing, chilling, blanching, fileting, cooking, and marination. It has been estimated that the processing of 1 t of raw seafood requires between 10 and 40 m^3^ of water (Arvanitoyannis and Kassaveti, 2008; Venugopal, 2021). Wastewater from the seafood industry typically contains high concentrations of complex organic substances, making it a major source of pollution (You et al., 2010; Chowdhury et al., 2010). In addition, its high salt content (in the range of 3%–15%) distinguishes it from other types of industrial wastewater and makes it difficult to treat using biological methods (Lefebvre and Moletta, 2006). Although physical and chemical technologies can be applied for treatment of industrial saline wastewater (e.g., membrane separation, physical adsorption, or electrodialysis), operational challenges such as membrane fouling, together with their high costs, still limit their widespread usage (Ferjani et al., 2005; Neilly et al., 2012). In addition, biological methods are often time-consuming and require substantial efforts to achieve the adaptation of salt-tolerant microorganisms (Ferjani et al., 2005). As a result, finding economically feasible and technologically efficient ways to treat saline wastewater from the seafood industry remains a challenge.

The use of microbial fuel cells (MFCs) has gained popularity as a promising and sustainable method for generating electrical energy from organic compounds present in wastewater, thus offsetting operating costs (Rossi and Logan, 2022; Boas et al., 2022). This technology not only helps in the elimination of organic pollutants but also generates electricity from liquid waste. Therefore, MFC application has become a significant concept in the field of wastewater treatment. However, few studies have examined whether and to what extent MFCs operated in continuous rather than batch mode can be used to remove organic compounds and produce energy from wastewater from seafood processing (Boas et al., 2022; Selvasembian et al., 2022).

In general, raw domestic and industrial wastewaters lack the necessary ionic conductivity to support high energy production in MFCs. As a result, they often require the addition of high concentration of inorganic salts, such as sodium chloride and phosphate buffer salts, to maintain an adequate solution ionic strength (Liu et al., 2005; Feng et al. 2008). Previous studies have shown that saline seafood wastewater has the potential to sustain power generation in MFCs due it its high ionic conductivity conditions (Logan and Rabaey, 2012; Lefebvre et al., 2012; Li et al., 2013a; Guo et al., 2021). However, these studies used inocula from domestic wastewater treatment plants (WWTPs) where microorganisms were not adapted to high saline conditions. The major disadvantage of the biological treatment of saline wastewater still is the inability of microorganisms to adapt to high saline levels (Lefebvre and Moletta 2006; Li et al., 2013a). Recent studies have shown that MFCs treating fish market wastewater inoculated with halophilic bacteria can achieve total chemical oxygen demand (COD) rates greater than 90% (Jamal and Pugazhendi, 2021). Yet, limited information is available on the use of MFCs to treat saline wastewater and produce energy using activated sludge from the seafood industry as inoculum (Xin et al., 2022).

The hydraulic retention time (HRT) is a critical factor in the design and operation of MFCs and can have a significant impact on organic matter (OM) removal and energy production. Previous studies showed that a higher HRT facilitates the efficiency of COD and OM removal in MFCs (Kim et al., 2015, 2016; Fazli et al., 2018). Other studies found that variations in HRT can influence the types and quantities of microorganisms in the anode, ultimately affecting the power output of MFCs (Sharma and Li, 2010; Sobieszuk et al., 2017). Additionally, the HRT can affect the level of shear stress in the anode chamber, which directly impacts the formation of biofilm on the anode surface (Lecuyer et al., 2011). Contrasting reports have been published on the impact of HRT on energy production. Whilst some studies found that a reduction in HRT is associated with an increased current production (Juang et al., 2012; Wei et al., 2012; Akman et al., 2013; Ge et al., 2013), others showed that a higher HRT enhanced current generation (Sharma and Li, 2010; Santos et al., 2017; Sobieszuk et al., 2017; Ye et al., 2020). Yet, it is still unknown how variations in the HRT may impact nutrient removal and current production in MFCs treating saline wastewater and operated in a continuous mode.

Exoelectrogenic prokaryotic (Bacteria and Archaea) and eukaryotic (mainly Fungi) organisms are the core elements of MFCs as they produce electrical currents directly from the oxidation of OM in the anode chamber and transfer electrons to a solid electrode through different mechanisms (Logan et al., 2019; Castellano-Hinojosa et al., 2022a). Therefore, the study of the characteristics of the anodic microbiome and its microbial interactions is critical to understand treatment performance and electricity production in MFCs. However, limited information is available on the abundance, diversity, and composition of prokaryotic and eukaryotic organisms in MFCs treating saline wastewater at different HRTs, although such information may help further refine this technology and select optimal operational parameters for achieving efficient OM removal and energy production.

In this study, we examined the impacts of three different HRTs on organic removal rate, energy production, and anodic microbial communities in an MFC treating saline wastewater and inoculated with activated sludge from the seafood industry. The abundance, diversity, and composition of prokaryotic and eukaryotic communities in the anode biofilm were studied and linked to physicochemical and electrochemical parameters.

## 2. **Materials and methods**

### 2.1. Design and operation of the MFC

A two-chambered H-cell type MFC reactor consisting of two methacrylate chambers (5 L for the anode and 4 L for the cathode), separated by a proton exchange membrane (Nafion N117, Chemours, Italy), was constructed (Supplementary Fig. S1). The anode (240 cm^2^ projected area) was of carbon fibers (6.35 mm thickness), and the cathode (17 cm^2^ projected area) was made of a copper bar. The distance between the electrodes was 60 cm, and the total length of the connected tube was 20 cm. The electrodes were positioned in the middle of each chamber, 5 cm from the wall (Supplementary Fig. S1). The anode and the cathode were connected on the outside of the MFC by means of a copper conductor cable for electron transport. The anode and cathode chambers were equipped with sensors for the continuous measurement of dissolved oxygen concentration, redox potential, pH, and temperature (Supplementary Fig. S1). The Nafion N117 membrane was pretreated by immersion in 5% NaCl solution for 12 hours to allow for membrane hydration and expansion, as recommended by the manufacturer.

The anode chamber was inoculated with 4 L of activated sludge from a fish- canning industry in Galicia (Northwest of Spain). This WWTP uses a conventional activated sludge (CAS) system for nutrient removal, operating at a moderate salinity level (12.76 NaCl/L), as described in full detail by Correa-Galeote et al. (2021a). The anode chamber was continuously fed from the top using a peristaltic pump with synthetic wastewater simulating fish-canning wastewater (Correa-Galeote et al., 2021b) with the following composition: CH_3_COONa 3H_2_O 5.6 g L^-1^ (2.5 g Ac^-^/L), NH_4_Cl 0.38 g L^-1^, MgSO_4_·7H_2_O 0.1 g L^-1^, K_2_HPO_4_ 0.085 g L^-1^, KCl 0.04 g L^-1^, KH_2_PO_4_ 0.03 g L^-1^, and NaCl 5.23 g L^-1^; pH was 7.1, and conductivity was 13.9 mS cm^-1^. The total organic carbon (TOC) and NaCl concentrations in the influent were adjusted to reach a final concentration of 1 and 10 g/L, respectively, as they are representative of common TOC and NaCl values in wastewater from fish-canning industries in Galicia (Northwest of Spain) (Correa-Galeaote et al., 2021a, b). Three consecutive HRTs were studied: 1 day (HRT1), 3 days (HRT3), and 6 days (HRT6) for 30 days, 21 days, and 21 days, respectively. The total duration of the experiment was 72 days. The organic loading rate (ORL) varied from 2.5 mg COD L^-1^ d^-1^, 0.8 mg COD L^-1^ d^-1^, and 0.4 mg COD L^-1^ d^-1^ at HRT1, HRT3, and HRT6, respectively. The catholyte was 50-mM phosphate buffer (PB; 4.58 g Na_2_HPO_4_, 2.45 g NaH_2_PO_4_; pH of 7.0; conductivity of 6.3 mS cm^-1^) and was renewed on a monthly basis (Rossi and Logan, 2020). The catholyte was continuously sparged with air to provide a dissolved oxygen (DO) level of 8.5 mg L^-1^, and both chambers were continuously mixed using a magnetic stirrer at 1,500 rpm. The MFC was operated at 20–22 °C in a controlled-temperature room.

### 2.2. Electrochemical measurements

The MFC was operated using an external resistance of 1,000 Ω during the experimental period. Current production was calculated applying Ohm’s law (I = V/R), where V is the measured voltage (volt), and R the external resistance (ohm). The power density, P (mW m^−2^), and the current density, j (mA m^−2^), in the anode were calculated according to the projected anode surface area (Logan, 2008), using Equations (1) and (2), respectively:

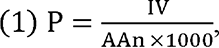

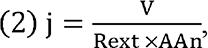

where I is the current (amp), V is the voltage (volt), Rext is the external resistance (ohm), and A_An_ is the projected anode area in m^2^.

The Coulombic efficiency (CE) was calculated as ‘current over time’ until the maximum theoretical current was achieved (Logan, 2008). The evaluated CE over time was calculated using Equation (1):

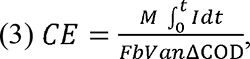

where M is the molecular weight of oxygen (32), F is Faraday’s constant (C/mol), b = 4 indicates the number of electrons exchanged per mole of oxygen (e-/mol), V_an_ is the volume of liquid in the anode compartment, and ΔCOD is the change in COD over time, ‘t’.

### 2.3. Physiochemical analyses

Physicochemical analysis was carried out twice per week over the experimental period, using samples (500 mL) collected from the influent and effluent. The COD was determined according to standard methods described by the APHA (APHA, 2012), and its removal % was calculated based on the concentrations in the influent and effluent. The organic removal rate (ORR) per day was calculated as the difference in the COD between the influent and influent divided by the HRT (1, 3, or 6 days). The concentrations of acetate (CH_3_C:COO−), ammonium (NH ^+^), nitrite (NO ^-^), and nitrate (NO ^-^) were analyzed using an ion chromatograph (Metrohm Ion Chromatograph, AG, Switzerland) in 0.22-µM-filtered samples. The removal % for OM and N were calculated as the difference in the concentration of CH_3_C:COO− and all inorganic N forms (NH ^+^ + NO ^-^ + NO ^-^), respectively, between the influent and effluent. Suspended solids in the effluent were determined according to the APHA (APHA, 2012). Conductivity in the influent and effluent was measured with a laboratory conductivity meter sensION+ pH3 (Hach Lange, Ames, USA). The values of pH, temperature, and redox potential were determined in the anode using sensors as described above.

### 2.4. DNA extraction and quantification of total bacterial, archaeal, and fungal communities

Biomass from the anode was collected after 7, 14, 21, 37, 44, 51, 58, 65, and 72 days of operation, corresponding to days 7, 14, and 21 for each of the three HRTs studied. Biomass form the original inoculum (day 0), the activated sludge from the seafood industry, was also used for DNA extraction. The anode biofilm was scraped from the anode using pre- sterilized tweezers and suspended in 20 mL of sterilized distilled water. The samples were then sonicated for 3 min and centrifuged at 13,000 rpm for 5 min. The pelleted biomass was kept at -20°C until use. The DNA was extracted using the FastDNA SPINK Kit for Soil (MP Biomedicals, Solon, OH, USA) and quantified using NanoDrop (Fisher Scientific, USA).

The abundances of total bacterial (16SB), archaeal (16SA), and fungal (18SF) communities in the anode biofilm were determined using a QuantStudio 3 Real-Time PCR system (ThermoFisher, USA) and the 16S rRNA and 18S rRNA genes as molecular markers. The PCR reaction mixtures and conditions, primers, and standards were described previously by Castellano-Hinojosa et al. (2018) and Maza-Márquez et al. (2019) and are presented in Supplementary Table S1. Standard curves were always linear (R^2^ > 0.998), with amplification efficiencies in the range of 92.7%–100%. The quality of the PCR amplification was confirmed through melting curve analyses and agarose gel electrophoreses, with no amplification detected in the no-template controls.

### 2.5. Analysis of prokaryotic and eukaryotic communities

Amplicon sequencing was conducted by Novogene Europe (Cambridge, UK) using the primer pairs Pro341F and Pro805R, which amplify the V3–V4 region of the 16S rRNA of prokaryotes (bacteria + archaea) (Takahashi et al., 2014), and the EUK1391 and EUKBr primers, which amplify the V9 region of the 18S rRNA gene of eukaryotes (Amaral-Zettler et al., 2009; Stoek et al., 2010), in an Illumina MiSeq sequencer. The sequence reads were analyzed in QIIME2 following the methods described in full detail by Castellano-Hinojosa et al. (2021). The final dataset consisted of an average of 58,654 and 25,654 sequences per sample for the prokaryotic and eukaryotic community, respectively. Raw sequence data were deposited in NCBI’s Sequence Read Archive under BioProject PRJNA967514.

Alpha [number of amplicon sequence variants (ASVs) as well as Shannon and Inverse Simpson indices] and beta diversity (analysis of unweighted UniFrac distances using non-metric multidimensional scaling, NMDS) analyses were carried out on log- normalized data to prevent errors caused by rarefaction (McMurdie and Holmes, 2014), using the R package “vegan” v. 2.5-2’ (Oksanen et al., 2016) and ’Phyloseq’ v. 1.24.0 (McMurdie and Holmes, 2013) in the R software v. 4.0.5 (http://www.rproject.org/). Community composition variations between HRTs and time points were evaluated by permutational analysis of variance (PERMANOVA). The DESeq2 package (Love et al., 2014) was employed to identify differentially abundant prokaryotic and eukaryotic ASVs between HRT1 *vs*. inoculum, HRT3 *vs.* HRT1, and HTR6 *vs.* HRT3, as described by Castellano-Hinojosa et al. (2022b). Significant Pearson correlations between differentially abundant taxa and voltage were identified using the “cor.test()” function in R.

### 2.6. Co-occurrence network analysis

The impacts of HRT1, HRT3, and HRT6 on prokaryotic and eukaryotic community organization and potential ecological interactions were assayed using co-occurrence networks, which were constructed as described in detail by Castellano-Hinojosa et al. (2022b), using Spearman correlations. Network properties such as the numbers of nodes and edges, mean degree, and density were inferred using the igraph package in R (Csárdi and Nepusz, 2006), and networks showing significant associations were constructed using the Cytoscape v. 3.9.0 software (Shannon et al., 2003).

### 2.7. Statistical analysis

The analysis of data was performed using the R software version 4.0.5 (http://www.rproject.org/). The Shapiro-Wilk test and Bartlett’s test were used to check if variables met the normality and homoscedasticity assumptions required for analysis of variance (ANOVA), respectively. One-way ANOVA comparisons of means and post-hoc (Tukey) tests were applied for comparisons between samples, and *p*-values ≤ 0.05 were considered significant. Redundancy analysis (RDA) was performed to assess the association between the total abundance of 16SB, 16SA, and 18SF communities and the physicochemical (N removal %, ORR, pH, temperature, redox potential, conductivity, and suspended solids) and electrochemical (voltage and CE) parameters, using the Canoco 5.0 software. Pearson’s correlation coefficients between vectors representing biotic and abiotic variables in the RDA plots were calculated.

## 3. **Results**

### 3.1. Effect of HRT on electrochemical parameters

The highest voltage and current density of 970 mV and 38.1 mA m^-2^, respectively, were obtained after 30 days of operation at HRT1 (Fig. 1A). Subsequently, the voltage and current densities gradually decreased to 639 mV and 26.3 mA m^-2^ on day 51 at HRT3, respectively, and to 578 mV and 24.1 mA m^-2^ by day 72 at HRT6, respectively (Fig. 1A). The power density produced by the MFC quickly increased during the first 30 days of operation and reached 39.2 mW m^-2^ on day 30 at HRT1, followed by a decrease to 17.1 mW m^-2^ by day 51 at HRT3 and to 13.6 mW m^-2^ by day 72 at HRT6 (Fig. 1B). Regardless of the HRT, a CE of approximately 3% was detected throughout the experimental period (Fig. 1B).

**Fig. 1.**
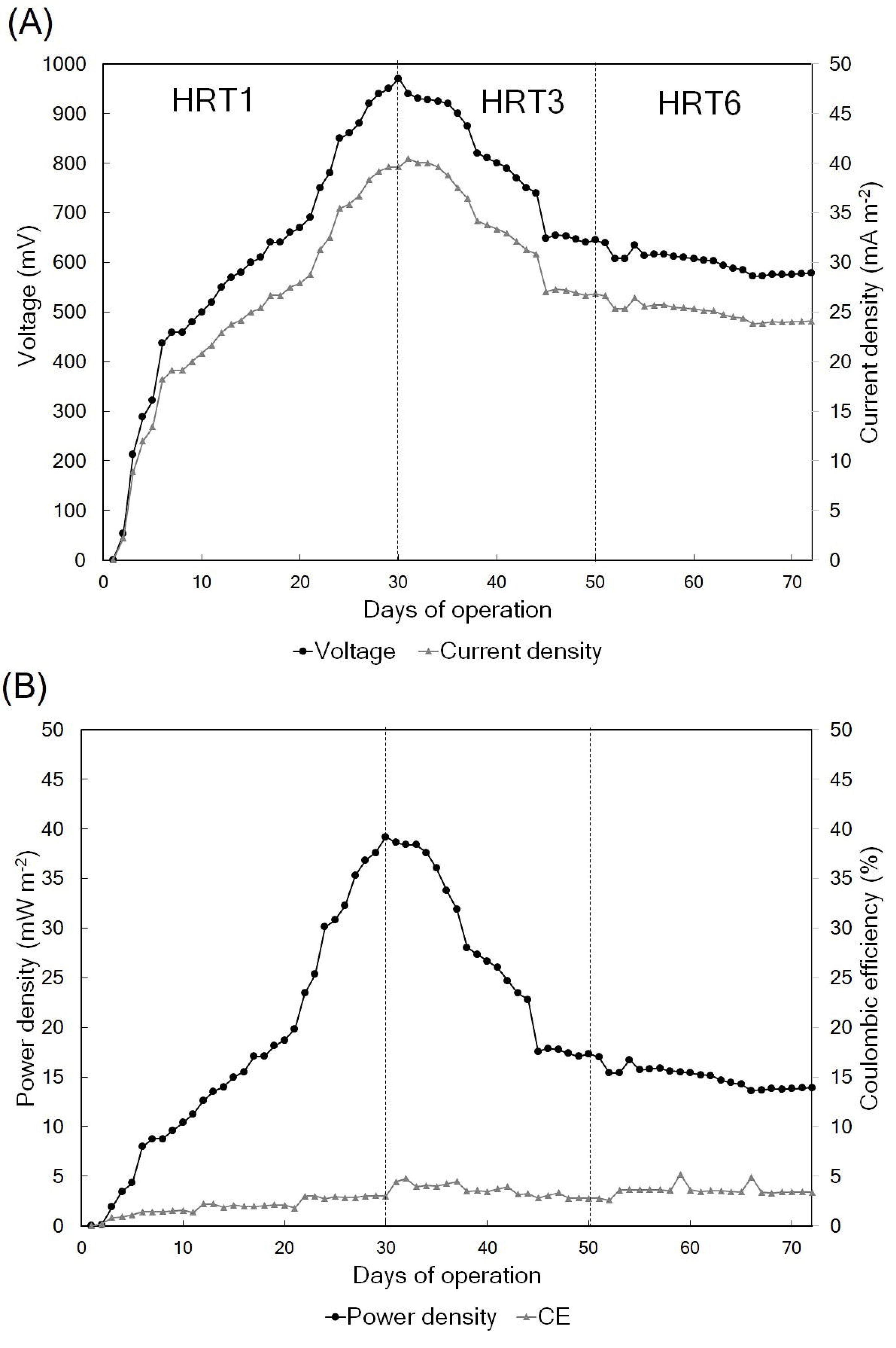
Voltage and current density (A) and power density and coulombic efficiency (B) generated during the experimental period. Three consecutive HRTs were examined: 1 day (HRT1), 3 days (HRT3), and 6 days (HRT6). CE, coulombic efficiency.

### 3.2. Effect of HRT on COD and OM removal and other physicochemical parameters

The COD (Fig. 2A) and OM (Fig. 2B) removal % were significantly greater at HRT6 compared to HRT3 and HRT1 and significantly greater at HRT3 compared to HRT1. However, the ORR significantly decreased with increased HRT, and was significantly greater at HRT1 compared to HRT3 and HRT1 (Fig. 2C).

**Fig. 2.**
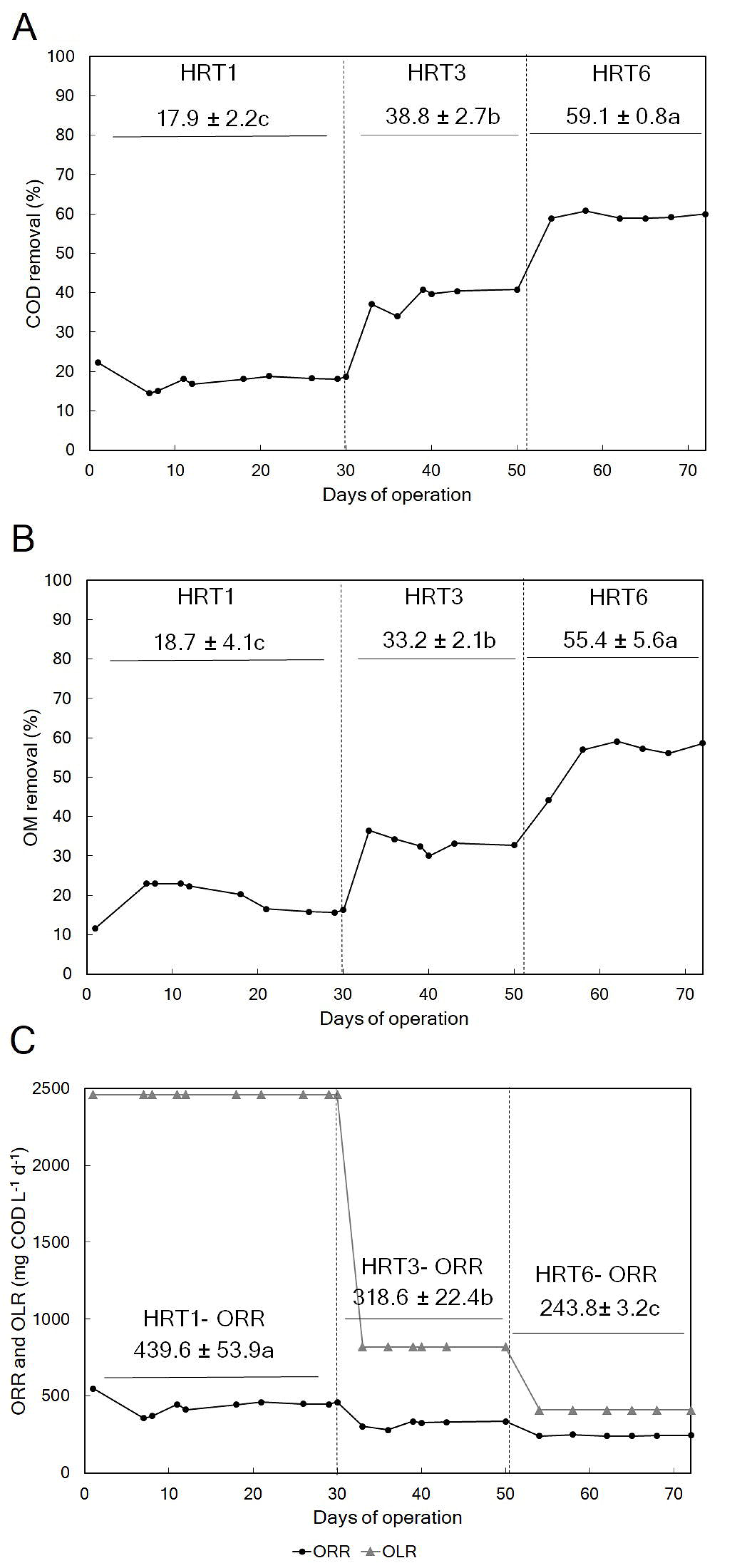
COD (A) and OM (B) removal %, and ORR and ORL (C) during the experimental period. Three consecutive HRTs were examined: 1 day (HRT1), 3 days (HRT3), and 6 days (HRT6). COD, chemical oxygen demand; OM, organic matter; ORR, organic removal rate. ORL, organic loading rate. Values are expressed as mean with standard deviation.

Regardless of the HRT, no significant differences in the values of pH (in the range of 7.8–8.2), temperature (in the range of 20.1–20.7°C), and redox potential (ranging from - 420 to -438 mV) were detected in the anode throughout the experimental period (Supplementary Table S2). No significant differences in the N removal % were detected among HRTs, with values in the range of 20.3%–28.5% throughout the 72 days of operation (Supplementary Table S2). The levels of suspended solids and conductivity in the effluent of the MFC varied between 32.6–49.6 mg L^-1^ and 11.2–12.3 mS cm^-1^ throughout the experimental period, but significant differences among HRTs were not detected (Supplementary Table S2).

### 3.3. Abundance of microbial communities and their relationships with electrochemical and physicochemical parameters

The total abundances of the 16SB (Fig. 3A) and 18SF (Fig. 3C) communities gradually and significantly increased at HRT1 until day 21 compared to the inoculum to remain unchanged until the end of the experiment at HRT3 and HRT6. The total abundance of the 16SA community was significantly greater at HRT3 and HRT6 compared to HRT1 (Fig. 3B). No significant differences in the total abundance of the archaeal 16S rRNA gene at HRT3 and HRT6 were detected throughout the experimental period (Fig. 3B).

**Fig. 3.**
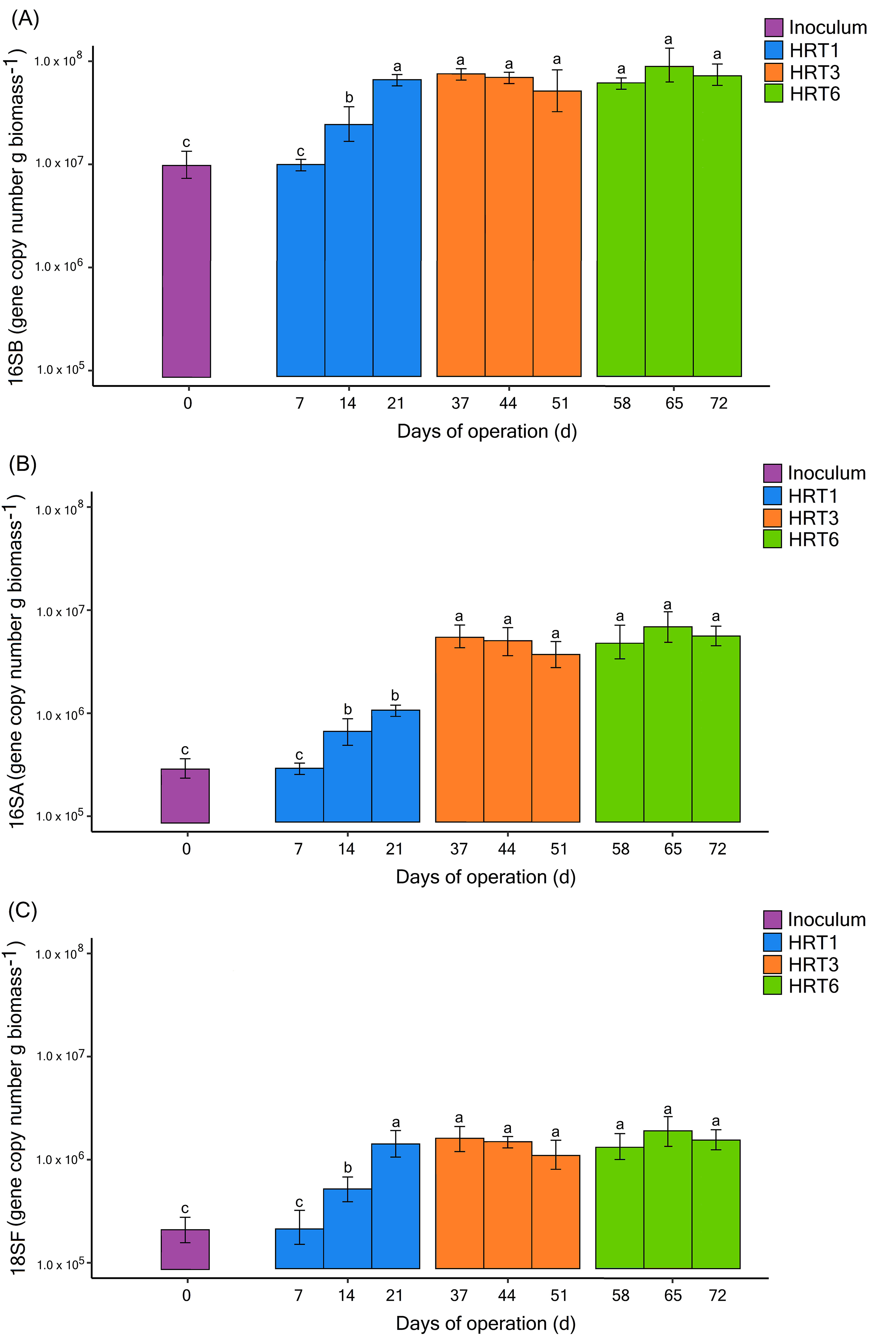
Total abundance of bacterial (16SB, A), archaeal (16SA; B), and fungal (18SF; C) communities at different time points during the experimental period. Three consecutive HRTs were run: 1 day (HRT1), 3 days (HRT3), and 6 days (HRT6). Different letters above the bars indicate significant differences between time points between HRTs according to one-way ANOVA with Tukey HSD test (*p* < 0.05; n = 5). Values are expressed as mean with standard error.

The results of the RDA, combined with the Pearson correlation coefficients, showed that the total abundance of 16SB was positively correlated (r > 0.73; *p* ≤ 0.01) with voltage, ORR, and CE. However, a strong negative correlation was found between the total abundance of 16SA and voltage, ORR, and CE (r < -0.85; *p* ≤ 0.01) (Supplementary Fig. S2; Supplementary Table S2). A significant positive correlation was found between the pH and the N removal % (r = 0.85; *p* ≤ 0.01) and between temperature and suspended solids (r = 0.75; *p* ≤ 0.05) (Supplementary Fig. S2; Supplementary Table S2). The abundance of 18SF was significantly and negatively correlated with the redox potential (r = -0.78; *p* ≤ 0.01). The values of conductivity significantly negatively correlated with the temperature and suspended solids (r < -0.75; *p* ≤ 0.05).

### 3.4. Diversity and composition of prokaryotic and eukaryotic communities

The use of HRT1, HRT3, and HRT6 had no significant effect on the number of ASVs compared to the inoculum for the prokaryotic community throughout the experimental period (Fig. 4A). However, significant increases in the values of the Shannon and Simpson indices were observed with time and greater HRTs (Fig. 4A). The application of any of the HRTs significantly increased the number of ASVs and the values of the Shannon and Simpson indices compared to the inoculum for the eukaryotic community (Fig. 4B). However, no significant differences in any of the alpha diversity indices were detected among HRTs for the eukaryotic community (Fig. 4B).

**Fig. 4.**
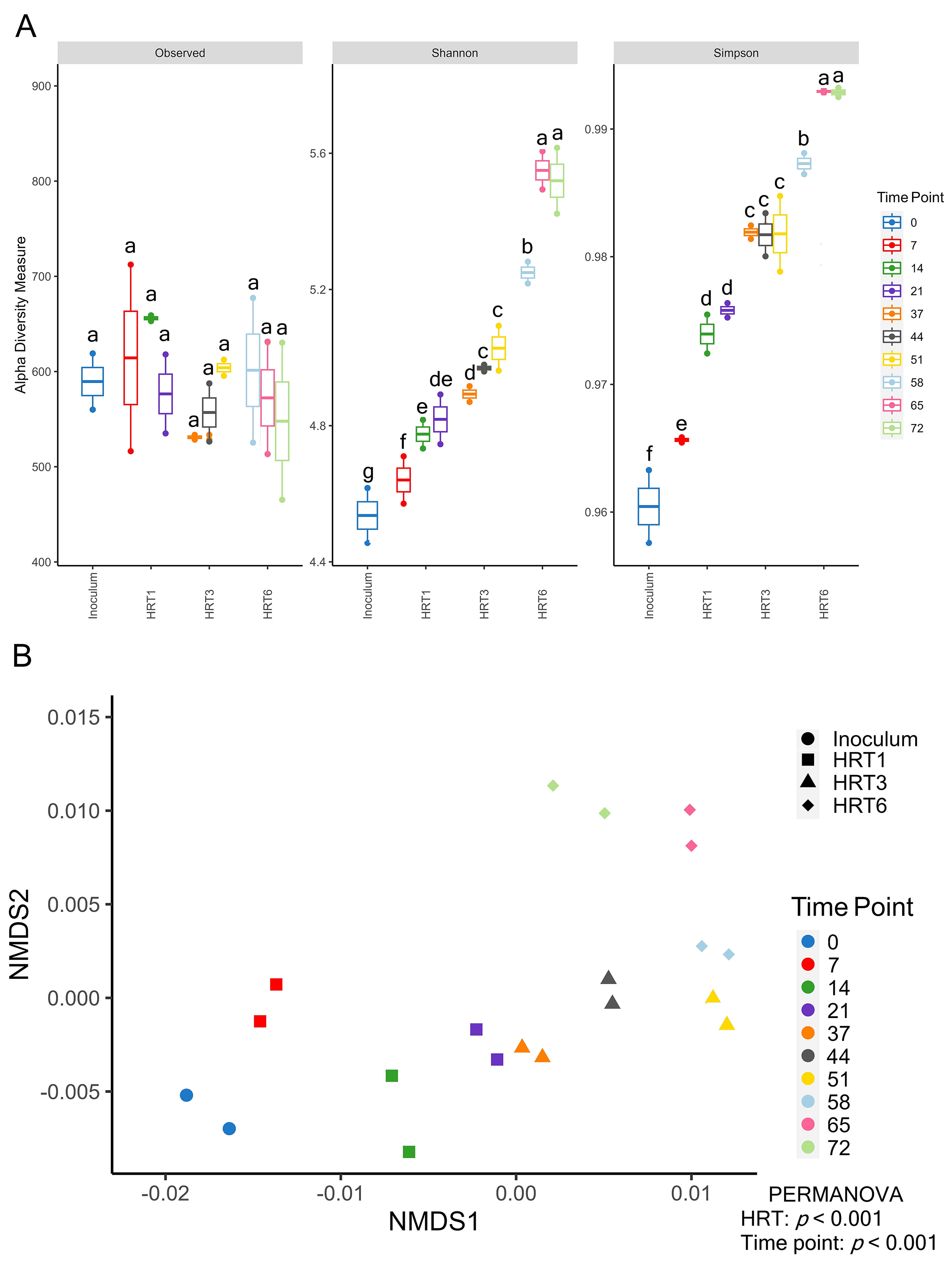
A. Number of ASVs, and values of Shannon and inverse Simpson diversity indices for the prokaryotic community at different time points during the experimental period. Different letters above the bars indicate significant differences between time points (Tukey’s HSD, *p* ≤ 0.05). Values are expressed as mean with standard error. B. Non-metric multidimensional scaling (NMDS) plots on unweighted UniFrac distances for the prokaryotic community at different time points during the experimental period. Three consecutive HRTs were run: 1 day (HRT1), 3 days (HRT3), and 6 days (HRT6). Differences in community composition between HRTs and time points were tested by permutational analysis of variance (PERMANOVA), and *p* values ≤ 0.01 were considered significant.

**Fig. 5.**
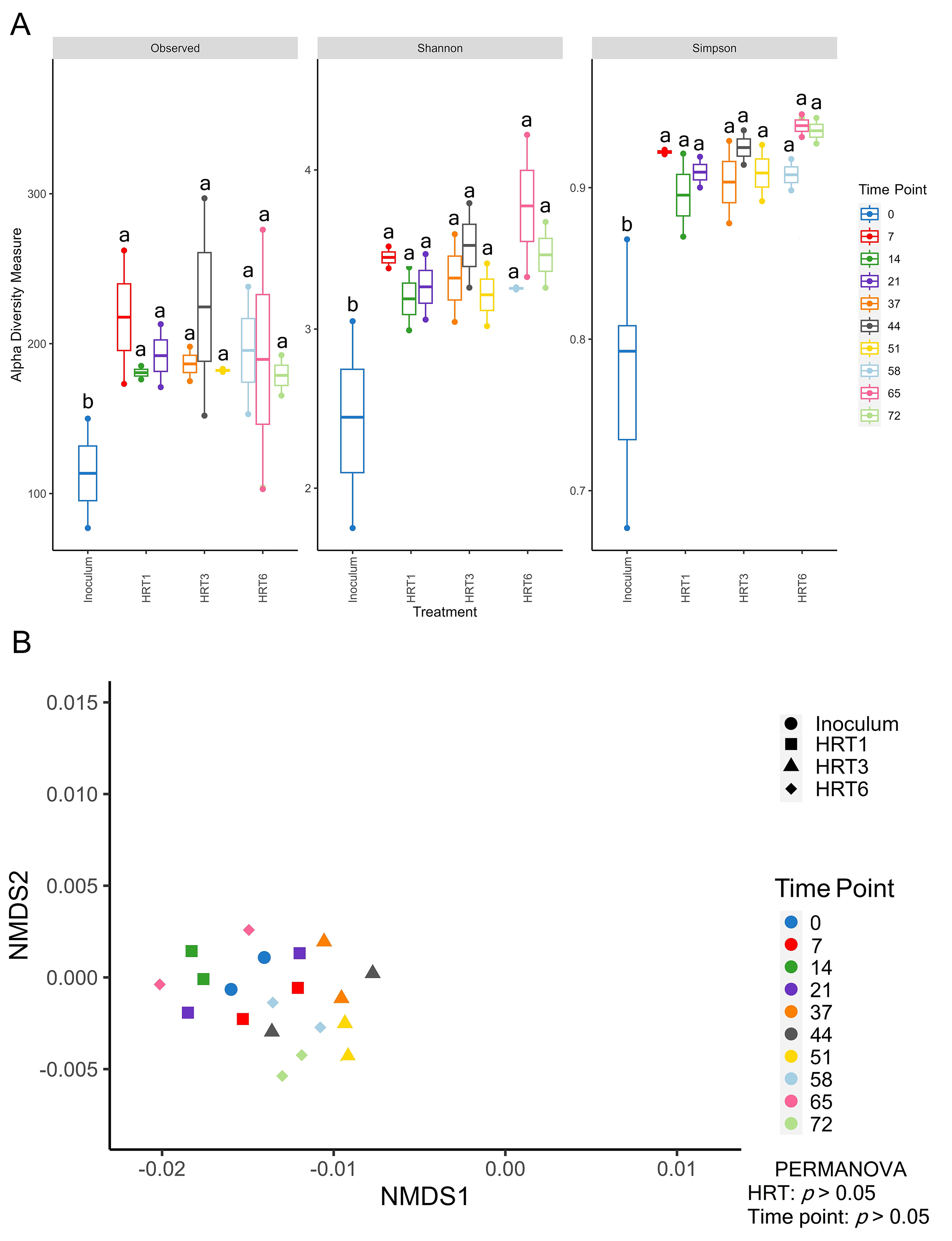
A. Number of ASVs, and values of Shannon and inverse Simpson diversity indices for the prokaryotic community at different time points during the experimental period. Different letters above the bars indicate significant differences between time points (Tukey’s HSD, *p* ≤ 0.05). Values are expressed as mean with standard error. B. Non-metric multidimensional scaling (NMDS) plots on unweighted UniFrac distances for the prokaryotic community at different time points during the experimental period. Three consecutive HRTs were run: 1 day (HRT1), 3 days (HRT3), and 6 days (HRT6). Differences in community composition between HRTs and time points were tested by permutational analysis of variance (PERMANOVA), and *p* values ≤ 0.01 were considered significant.

The NMDS analyses on unweighted UniFrac distances, together with the PERMANOVA, showed significant differences in the composition of the prokaryotic community between HRTs and time points (*p* ≤ 0.001). No significant differences in beta diversity were detected between HRTs and time points (*p* ≥ 0.05) for the eukaryotic community (Fig. 4B).

On average, Pseudomonadota (40.99 %), Chloroflexota (18.5 %), Actinomycetota (10.1%), and Bacillota (9.7 %) were the most abundant prokaryotic phyla across all HRTs and time points (Supplementary Fig. S3A). Rhodospirillaceaeae, Anaerolineaceae, and Gemmatimonadaceae were the dominant prokaryotic families in the anodic microbiome (Supplementary Fig. S3B). The eukaryotic community was mainly formed by the phyla Ascomycota (50.5%) and Basidiomycota (26.8%) (Supplementary Fig. S4A), as well as the families Saccharomycetaceae (42.9%) and Basidiomycota (11.5%) (Supplementary Fig. S4B).

### 3.5. Differentially abundant prokaryotic and eukaryotic taxa and their relationships with physicochemical and electrochemical parameters

Prokaryotic ASVs significantly enriched and depleted between HRT1 and the inoculum, between HRT3 and HRT1, and between HRT6 and HRT3 were identified at the genus taxonomic level (Fig. 6). The application of HRT1 caused significant increases in the relative abundances of ASVs belonging to the genera *Acholeplasma*, *Arcobacter*, *Candidatus* Mirothrix, *Geobacter*, *Crostidium*, and *Shewanella* compared to the inoculum (Fig. 6A). The use of HRT3 mainly enriched ASVs belonging to the archaeal genus *Methanosarcina* compared to HRT1, whereas it depleted those belonging to 19 bacterial genera including *Clostridium*, *Geobacter*, *Candidatus* Mirothrix, and *Shewanella* (Fig. 6B). The application of HRT6 significantly enriched and depleted ASVs belonging to more than 40 different bacterial genera compared to HRT3 and particularly favored the proliferation of taxa from the archaeal genus *Methanosarcina* (Fig. 6C). No differentially abundant eukaryotic taxa were detected between the inoculum and HRT1 and among the HRTs.

**Fig. 6.**
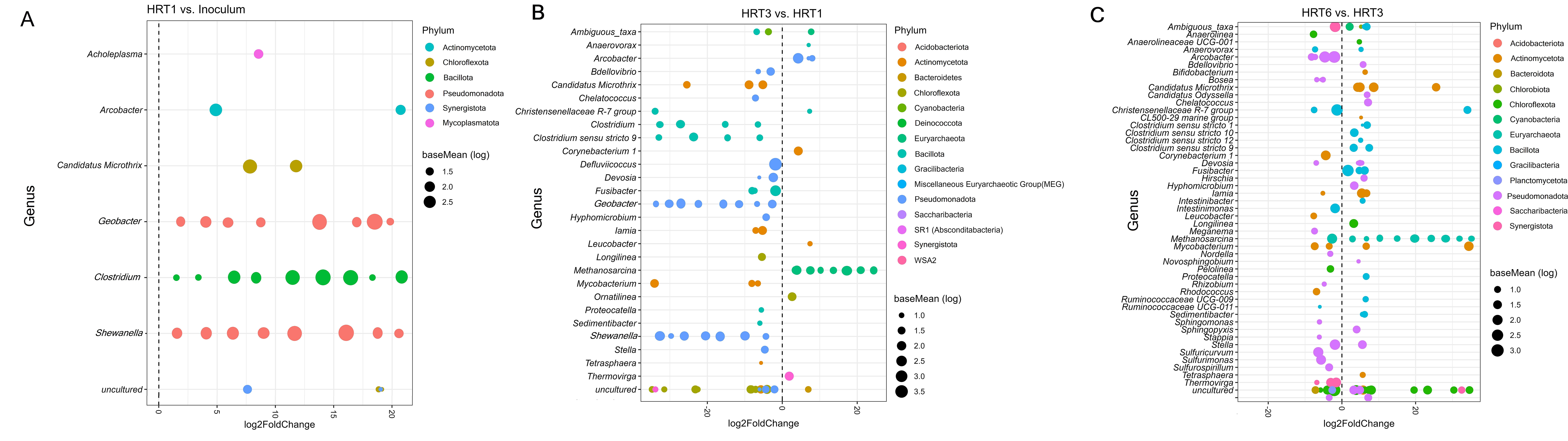
Differential abundance prokaryotic ASVs at the genus taxonomic level between the inoculum and HRT1, HRT1 and HRT3, and HRT3 and HRT6. Three consecutive HRTs were run: 1 day (HRT1), 3 days (HRT3), and 6 days (HRT6). Each colored dot represents an ASV that was identified by DESeq2 analysis as significantly differentially abundant between treated and non-treated soils (*p* ≤ 0.05).

Among the differentially abundant prokaryotic taxa identified in Fig. 6, a total of 25 bacterial genera and 1 archaeal genus were significantly correlated with voltage (Fig. 7). The genera *Clostridium*, *Geobacter*, and *Shewanella* showed the strongest correlations with current production (r > 0.80), whereas others such as *Candidatus* Microthix, *Longilinea*, *Ornatilinea*, *Anaerolinea*, *Clostridium sensu stricto 1*, *Bdellovibrio*, and *Corynebacterium* showed significant positive correlations with voltage in the range of 0.55–0.68 (Fig. 7). The bacterial genera *Proteocatella*, *Candidatus* Odyssella, and *Slufurospirillum*, as well as the archaeal genus *Methanosarcina*, were significantly correlated with voltage (Fig. 7).

**Fig. 7.**
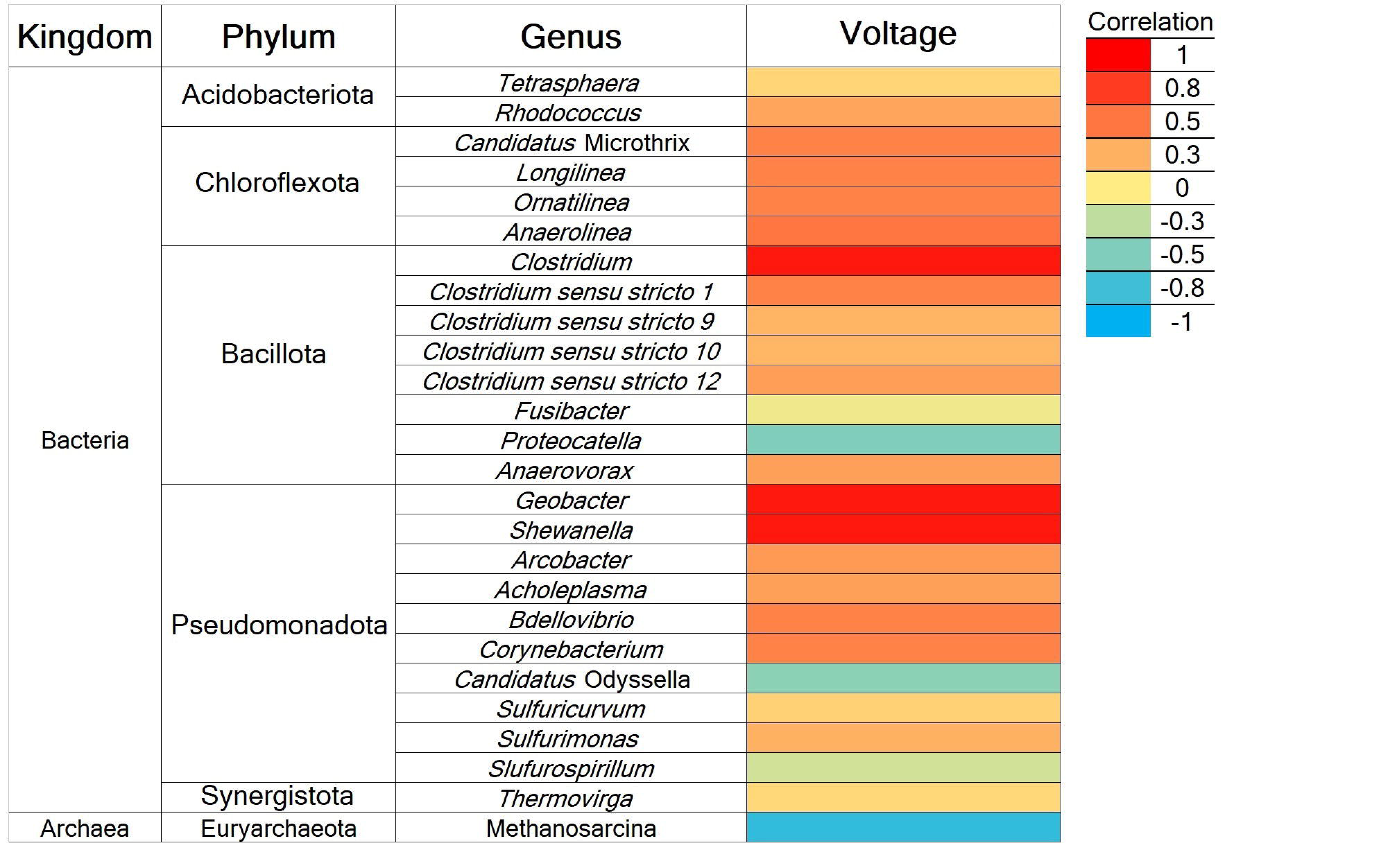
Significant Pearson correlations between the differentially abundant prokaryotic taxa detected in Fig. 6 and voltage (*p* ≤ 0.05). The shading from blue to red represents low- to high-positive Pearson correlation coefficients.

### 3.6. Effect of HRT on microbial network complexity

The co-occurrence networks of prokaryotic communities (Supplementary Fig. S5) were constructed to explore the co-occurrence patterns of anodic microbes throughout the experimental period at different HRTs. The density and numbers of edges and nodes from the prokaryotic networks gradually and significantly increased with time and greater HRTs, and network complexity was highest at HRT6 (*p* ≤ 0.05; Supplementary Fig. S5; Supplementary Table S4). No significant differences in network complexity were detected for the eukaryotic community throughout the experimental period (data not shown).

## 4. **Discussion**

We found that the HRT determines variations in organic removal rate, voltage, and the anodic microbiome of an MFC inoculated with activated sludge from a seafood industry and operated in a continuous mode. Decreases in the efficiency of removal of organic compounds (ORR) and increases in the abundance of archaeal communities with increased HRT was related to limited CE at greater HRT. The use of different HRTs (1, 3, and 6 days) determined significant variations in the abundances of bacterial and archaeal communities, and the diversity, composition, and network complexity of prokaryotic communities, which were strongly linked to changes in electrochemical and physicochemical parameters. For example, higher ORR and energy production at HRT1 was tightly linked to increased and decreased absolute abundances of bacterial and archaeal communities, respectively. The eukaryotic community was less responsive to changes in HRT, had no impact on current production but contributed to the removal of COD and OM. We showed that the use of HRT1 favored increases in the relative abundance of a diverse group of known electroactive microorganisms, including *Geobacter*, *Shewanella*, *Arcobacter*, and *Clostridium*, compared to HRT3 and HRT6. The proliferation of archaeal communities (mainly those belonging to the genus *Methanosarcina*) was related to decreased energy production and ORR at HRT3 and HRT6 compared to HRT1, likely due to strong competition between exoelectrogenic microorganisms and methanogens for electrons. Our results suggest that the anodic community of the MFC had the potential to produce electricity while efficiently removing high concentrations of COD and OM. These results highlight the feasibility of MFC systems to treat influents with moderate salinity and high TOC, such as wastewater from the fish-canning industry.

The voltage, current density, and power density were increased at lower HRTs, whereas CE did not vary among HRTs. Decreases in energy production at a longer retention time have been previously observed in MFCs operated in a continuous flow and fed with domestic wastewater (Liu et al., 2008; Sharma and Li, 2010). This is thought to be due to decreases in cell metabolism and/or the fouling of the electrodes caused by cell decay or death at a longer HRT (Walter et al., 2022). The gradual increases in voltage during the first 30 days of operation at HRT1 in our study could be due to a rapid and efficient use of organic compounds (as revealed by changes in ORR) and the subsequent electron transfer by the exoelectrogenic microorganisms at lower HRTs. The lower HRT favored increases in the abundance of bacterial communities which positively influenced CE and voltage. These results suggest that exoelectrogenic bacteria may have a greater ability to colonize anodes than other microbial communities, which favors rapid electron transfer at HRT1. Decreases in the efficiency of removal of organic compounds (ORR) with increased HRT can explain the decreases in current production at HRT3 and HRT6 compared to HRT1. In addition, the proliferation of archaeal communities at HRT3 and HRT6 was related to the decreased current generation. This is in line with other studies showing that the proliferation of electroactive methanogenic archaea (e.g., methane production; Karuppiah et al., 2018) and/or non-electroactive microorganisms at greater HRTs may limit energy production through a competition with exoelectrogenic microbes for electrons (Li et al., 2016; Karuppiah et al., 2018; Godain et al., 2021).

Regardless of the HRT, our results show the activated sludge from the fish-canning industry can be a good inoculum for energy generation in MFCs. Previous studies have shown that wastewater from the seafood industry may be more enriched with electroactive microorganisms compared to activated sludge from domestic WWTPs as substrate salinity appears to favor the growth of electroactive microorganisms (Guo et al., 2021; Xin et al., 2022). This is supported by the detection of greater values of current density and power density in our study compared to other MFCs inoculated with activated sludge from domestic wastewater treatment plants and fed with seafood wastewater (You et al., 2010; Sukkasem and Laehlah, 2013; Tremouli et al., 2017; Jayashree et al., 2016). The use of saline influents increases ionic conductivity and decreases the resistance of anaerobic electrolytes, which enhances ion transfer in the anode chamber and can increase the power output of MFCs (Velasquez-Orta et al., 2011; Guo et al., 2021; Xin et al., 2022). Although electroactive fungi can contribute to energy production in MFCs (Logan et al., 2019; Castellano-Hinojosa et al., 2022a), they had no significant impact on the electrochemical parameters in our study.

As expected, when the HRT increased from 1 day to 3 and 6 days with the same influent COD concentration (2.4 gCOD/L), the COD removal % increased. These results agree with those found in other studies, where higher HRTs increased COD removal % in MFCs treating domestic wastewater (Li et al., 2013b; Kim et al., 2015, 2016; Fazli et al., 2018). However, we found that the ORR, which is the amount of COD removed per liter per day, decreased when the HRT was increased. These results shows that the efficiency of the COD removal was lower at higher HRT and lower OLR which resulted in a lower power density production. This can be explained by the lower concentration of substrate that can be utilized by electrogenic bacteria at lower OLR which resulted in lower ORR and power generation. Our results agree with those of Sharma and Li (2010) who reported reduced power density when the OLR was decreased from 6.5 g L^-1^d^-1^ to 1.4 g L^-1^d^-1^ operating at 23 h of HRT. They found an optimum OLR within the range of 2.35-3.44 g L^-1^d^-1^. Similarly, Liu et al. (2018) obtained higher power density when the HRT was increased from 4.1 h to 11.3 h and the OLR was decreased from 5.18 g L^-1^d^-1^ to 2.18 g L^-1^d^-1^ with a constant increased in the acetate removal %. However, when the HRT was increased from 11.3 h to 16 h (OLR was decreased from 2.18 g L^-1^d^-1^ to 1.5 g L^-1^d^-1^) the power density decreased and the acetate removal % increased less proportionally which showed that the efficiency of organic matter removal and energy production decreased from an optimum OLR of around 2.18 g L^-1^d^-1^ (Liu et al. 2008). A previous study using a halophilic consortium as inoculum and fish market wastewater reported a COD removal of 84% (initial concentration of 1.21 g COD/L) at an HRT of 20 days (Jamal and Pugazhendi, 2021). Here, a COD removal % of approximately 60% was obtained at an HRT of only 6 days with twice the initial COD (2.46 g COD/L). Our results suggest that the microbial communities in the activated sludge from the seafood industry had not only the potential to colonize anodes and produce energy but also to efficiently remove organic compounds.

The HRT determined the variations in the diversity and composition of the prokaryotic community, which critically impacted voltage. Increases in the HRT increased the diversity of the prokaryotic community without altering the number of total ASVs. This was further supported by detecting numerous significantly enriched prokaryotic genera at HRT3 and HRT6 compared to HRT1. Microbial community analysis showed that ASVs belonging to known electroactive bacterial genera were enriched at a lower HRT. For example, *Geobacter*, *Shewanella*, *Candidatus* Microthix, *Longilinea*, *Ornatilinea*, *Anaerolinea*, *Clostridium sensu stricto 1*, *Bdellovibrio*, and *Corynebacterium* were significantly positively correlated with voltage at HRT1. All these genera contain species that can transfer electrons to anodes in MFCs (Koch and Harnisch, 2016; Logan et al., 2019; Lovley and Holmes, 2022). Again, these results suggest that activated sludge from wastewater of the fish-canning industry can be a good source of electroactive microorganisms for MFCs. Increases in the HRT enriched methanogens such as *Methanosarcina* sp. and other bacterial genera, including *Proteocatella*, *Candidatus* Odyssella, and *Sulfurospirillum*. Exoelectrogenic archaea, such as *Methanosarcina*, compete for electrons with other groups of electroactive microorganisms in MFCs (Logan et al., 2019; Ramanaiah et al., 2021), thus reducing electron transfer and, ultimately, limiting current production. *Sulfurospirillum* sp. uses electrons for biohydrogen production (Kruse et al., 2018). To our knowledge, this is the first report on the potential role of *Proteocatella* and *Candidatus* Odyssella on current generation in MFCs.

Increases in the HRT increased prokaryotic network complexity throughout the experimental period. Microbial communities establish complex ecological networks, which can be important for maintaining the stability of the microbiome in response to external stresses (Banerjee et al., 2019). Increases in network complexity and alpha diversity, and alterations in the prokaryotic community composition, suggest that the increases in the HRT induced an increase in some microbial functions. Whether and to what extent these potential changes in the functionality of the anodic microbiome can be related to the performance and current production of MFCs should be explored in future studies by using tools such as shotgun metagenome sequencing.

In this study, the eukaryotic community did not respond to variations in HRT and played no significant role in energy production. Nevertheless, the diversity of the eukaryotic community increased after inoculation of the MFC, suggesting that some eukaryotic groups proliferated in the anode. Because fungi belonging to the phyla Ascomycota and Basidiomycota dominated the eukaryotic community, they might have contributed to COD and OM removal. It should be noted that studies of the absolute abundance and diversity of prokaryotic and eukaryotic communities does not distinguish between exoelectrogenic and non-exoelectrogenic microorganisms as both can coexist in

MFCs and can simultaneously impact system performance (e.g., COD and OM removal) and energy production (Asensio et al., 2017; Logan et al., 2019). Yet, it remains challenging to distinguish between the abundances of electroactive and non-electroactive microorganisms in anode biofilm in MFCs.

## 5. **Conclusions**

There is an increased interest in the valorization of industrial effluents using the MFC technology, but limited knowledge is available on those from the fish-canning industry in MFCs operated in a continuous mode. This study showed a close linkage between variations in the HRT and changes in physicochemical and electrochemical parameters and the abundances of bacterial and archaeal communities. Decreases in the efficiency of removal of organic compounds and increases in the abundance of archaeal communities with increased HRT was related to limited energy production at greater HRT. Increased efficiency of removal of organic compounds and energy production was detected at HRT of 1 day compared to HRT of 3 and 6 days. The HRT determined the variations in the relative abundances of specific bacterial and archaeal taxa contributing to increases and decreases in voltage. A diverse group of known electroactive bacterial genera contributing to increased voltage was identified, whereas mainly methanogens of the genus *Methanosarcina* were related to decreased current generation. Future studies should explore the impacts of different HRTs and OLR on the performance and energy production using different inoculum types. Microbial studies on the impact of HRT and OLR on the functionality of the anodic microbiome may increase our understanding of microbe- electrode interactions in MFCs treating industrial effluents.

## Data availability

Data will be made available on request.

## Acknowledgments

This work was supported by the P20-00079 grant of Junta de Andalucía of Proyectos I+D+i 2020. ACH is recipient of a Marie Skłodowska-Curie Postdoctoral European Fellowship (101108081) from HORIZON-MSCA-2022-PF-01 (Horizon Europe, 2022). We thank the research group of Dr. Anuska Mosquera-Corral for providing the activated sludge from the seafood industry for this study.

