## Supplementary Material for "Hydraulic retention time drives changes in energy production and the anodic microbiome of a microbial fuel cell (MFC)"

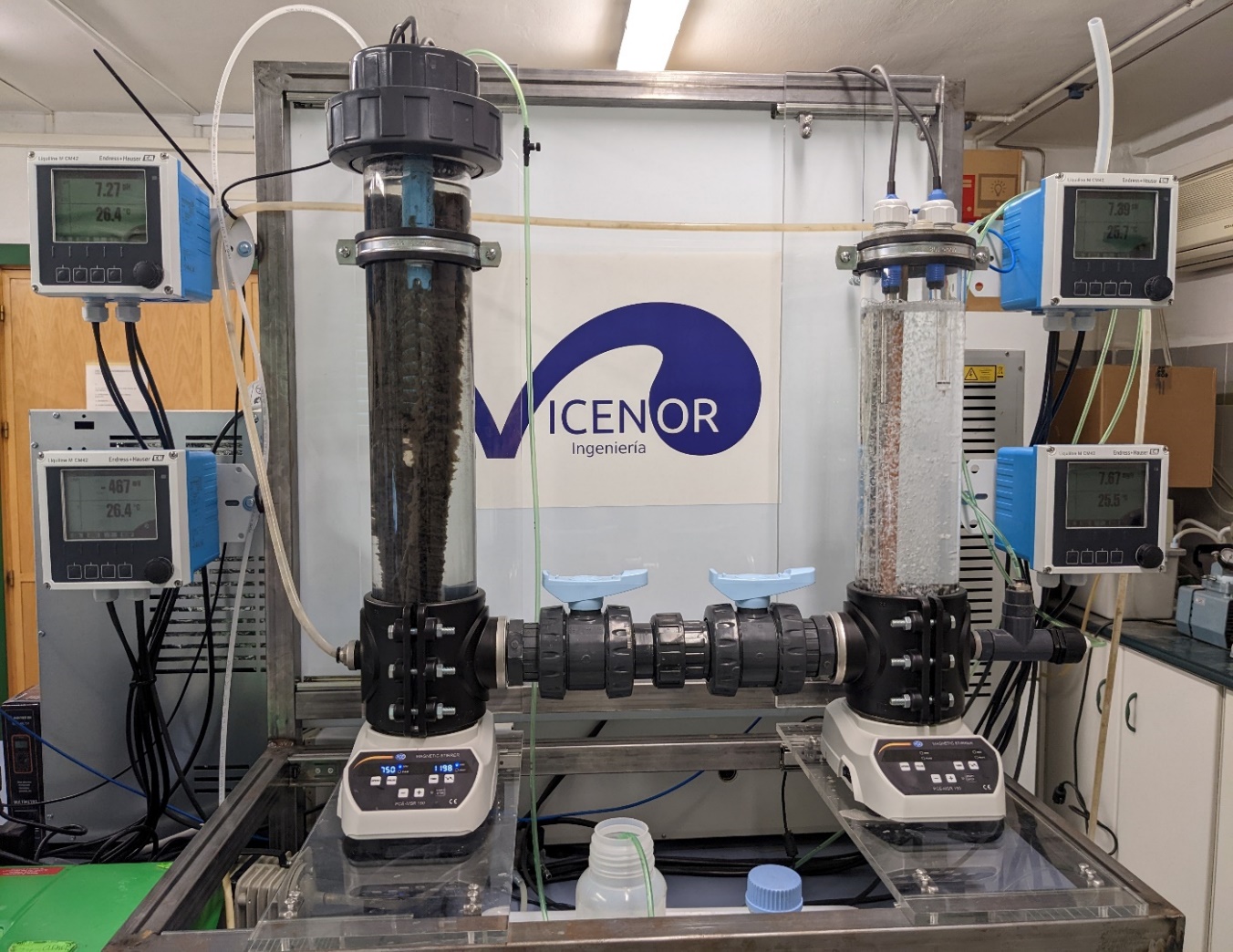

**DO sensor**

**Magnetic mixer**

**Carbon fibers**

**pH sensor**

**pH sensor**

**Redox sensor**

**PEM**

**Cathode chamber**

**Anode chamber**

**Supplementary Fig. S1.** A picture of the microbial fuel cell (MFC) used in this study. PEM, proton exchange membrane; T, temperature; DO, dissolved oxygen.

**Supplementary Table S1.** Primers used for quantification of total abundance of Bacteria, Archaea, and Fungi (16SB, 16SA, and 18SF, respectively) by qPCR (a). Bacterial, archaeal, and fungal strains used for generation of qPCR standards are also included. qPCR conditions for the quantification of each of the target genes (b).

a.

| Primer | Primer sequence (5´-3´) | Target gene | Strains | | Reference | |
| --- | --- | --- | --- | --- | --- | --- |
| 341F | CCTACGGGAGGCAGCAG | 16S rRNA Bacteria | | *Pseudomonas putida* NCB957 | | Muyzer et al. (1993) |
| 534R | ATTACCGCGGCTGCTGG |  |  |  |  |  |
| ARCH915F | AGGAATTGGCGGGGGAGCAC | 16S rRNA Archaea | | Genomic clone 29i4 | | Yu et al. (2008) |
| UNI-b-revR | GACGGGCGGTGTGTRCAA |  |  |  |  |  |
| FungiQuant-F | GSWCTATCCCCAKCACGA | 18S rRNA Fungi | | *Candida albicans* ATCC 10231 | | Liu et al. (2012) |
| FungiQuant-R | GGRAAACTCACCAGGTCCAG |  |  |  |  |  |

b.

|  | 16SB | 16SA | 18SF |
| --- | --- | --- | --- |
| Stage 1: 1 cycle | 3 min at 95 ºC | 7 min at 95 ºC | 3 min at 95 ºC |
| Stage 2: 40 cycles | 15s at 95 ºC | 30s at 95 ºC | 30s at 94 ºC |
|  | 30s at 60ºC | 30s at 65 ºC | 30s at 62 ºC |
|  | 30s at 72 ºC | 30s at 72 ºC | 45s at 72 ºC |
| Stage 3: 1 cycle | 10 min at 72 ºC | 10 min at 72 ºC | 10 min at 72 ºC |

**Supplementary Table S2.** Changes in the pH, temperature, and redox potential in the anode, conductivity and suspended solids in the effluent, and N removal efficiency (%) during the experimental period. Three consecutive HRTs were examined: 1 day (HRT1), 3 days (HRT3), and 6 days (HRT6). Values are expressed as mean with standard error. For each column, values followed by the same letter are not statistically different according to one-way ANOVA with Tukey HSD test (*p* < 0.05).

| HRT | Day of operation | pH anode | Temperature (ºC) anode | Redox potential anode (mV) | Conductivity effluent (mS cm^-1^) | Suspended solids effluent (mg L^-1^) | %N removal |
| --- | --- | --- | --- | --- | --- | --- | --- |
| HRT1 | 2 | 7.8 ± 0.2a | 20.1 ± 0.3a | -423 ± 5a | 11.2 ± 0.2a | 32.6 ± 1.1a | 21.6 ± 2.5a |
|  | 5 | 7.9 ± 0.1a | 20.2 ± 0.2a | -421 ± 6a | 11.3 ± 0.3a | 33.8 ± 1.9a | 22.1 ± 2.0a |
|  | 7 | 7.9 ± 0.2a | 20.1 ± 0.2a | -433 ± 5a | 11.5 ± 0.3a | 40.5 ± 3.1a | 21.8 ± 1.5a |
|  | 11 | 8.0 ± 0.2a | 20.0 ± 0.1a | -433 ± 7a | 11.5 ± 0.2a | 38.4 ± 1.8a | 23.2 ± 2.3a |
|  | 14 | 7.9 ± 0.1a | 20.3 ± 0.2a | -435 ± 5a | 12.1 ± 0.3a | 39.6 ± 1.9a | 28.5 ± 1.9a |
|  | 21 | 7.9 ± 0.2a | 20.2 ± 0.1a | -436 ± 8a | 12.0 ± 0.2a | 40.2 ± 2.7a | 25.4 ± 1.7a |
|  | 30 | 7.9 ± 0.2a | 20.5 ± 0.2a | -436 ± 5a | 11.8 ± 0.3a | 40.9 ± 1.9a | 27.2 ± 3.1a |
| HRT3 | 32 | 8.0 ± 0.2a | 20.5 ± 0.2a | -420 ± 6a | 12.3 ± 0.3a | 50.1 ± 2.6a | 25.4 ± 3.5a |
|  | 37 | 8.0 ± 0.2a | 20.4 ± 0.3a | -422 ± 5a | 11.5 ± 0.2a | 50.5 ± 2.2a | 26.2 ± 2.5a |
|  | 40 | 8.0 ± 0.1a | 20.3 ± 0.2a | -430 ± 5a | 11.9 ± 0.3a | 40.5 ± 3.3a | 22.4 ± 2.2a |
|  | 43 | 7.9 ± 0.2a | 20.6 ± 0.1a | -436 ± 10a | 11.8 ± 0.4a | 39.5 ± 1.4a | 25.7 ± 2.6a |
|  | 48 | 7.8 ± 0.3a | 20.5 ± 0.2a | -436 ± 5a | 12.1 ± 0.3a | 45.2 ± 1.9a | 23.9 ± 1.8a |
|  | 51 | 8.0 ± 0.2a | 20.4 ± 0.3a | -436 ± 7a | 12.0 ± 0.2a | 40.9 ± 1.8a | 26.5 ± 2.9a |
| HRT6 | 58 | 8.2 ± 0.2a | 20.6 ± 0.2a | -435 ± 5a | 11.6 ± 0.3a | 49.6 ± 2.0a | 20.3 ± 3.0a |
|  | 60 | 8.1 ± 0.2a | 20.5 ± 0.1a | -435 ± 9a | 11.5 ± 0.2a | 47.2 ± 2.6a | 26.5 ± 2.2a |
|  | 65 | 8.0 ± 0.1a | 20.6 ± 0.2a | -438 ± 5a | 11.4 ± 0.3a | 40.5 ± 3.6a | 22.2 ± 1.8a |
|  | 67 | 8.1 ± 0.2a | 20.4 ± 0.3a | -438 ± 6a | 11.7 ± 0.4a | 38.6 ± 2.9a | 25.4 ± 3.6a |
|  | 69 | 7.9 ± 0.1a | 20.5 ± 0.2a | -438 ± 5a | 12.0 ± 0.3a | 32.6 ± 3.1a | 23.2 ± 3.5a |
|  | 72 | 7.8 ± 0.2a | 20.7 ± 0.2a | -438 ± 8a | 12.0 ± 0.2a | 38.9 ± 2.8a | 28.5 ± 1.9a |

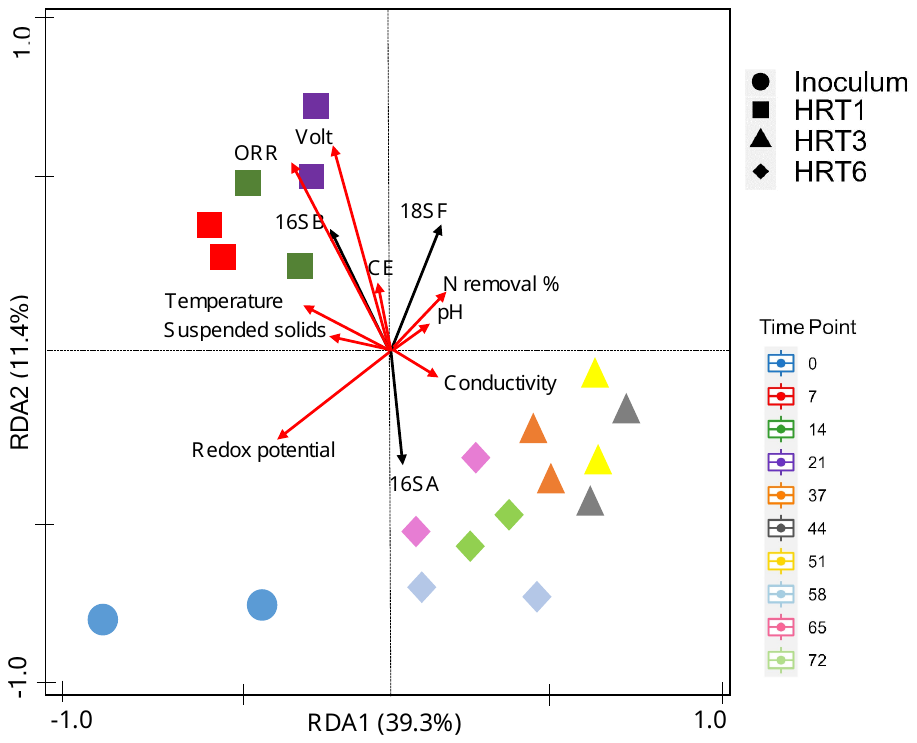

**Supplementary Fig. S2.** RDA plots of correlation between physicochemical and electrochemical parameters and abundance of total bacterial (16SB), archaeal (16SA), and fungal (18SF) communities during the experimental period. Samples from the inoculum and each of the three hydraulic retention times (HRT) of 1 day (HRT1), 3 days (HRT3), and 6 days (HRT6) tested were included. Red solid arrows indicate physicochemical and electrochemical factors, and blue arrows indicate total abundances of target genes.

**Supplementary Table S3.** Pearson’s product-moment correlation coefficients between the vectors displayed in Supplementary Fig. S2, which represent the total abundance of bacterial (16SB) and archaeal (16SA), and fungal (18SF) microbial communities and physicochemical and electrochemical parameters (%OM, %COD, and %N removal, pH, DO, redox potential, and voltage production) measured during the experimental period. Correlations between the gene abundances and abiotic variables are presented. Significant correlations (*p* ≤ 0.05) are boldfaced. ORR, organic removal rate; N, nitrogen; Volt, voltage; CE, coulombic efficiency.

|  | 16SB | 16SA | 18SF | ORR | N removal % | Redox potential | Voltage production | CE | pH | Temperature | Conductivity | Suspended solids |
| --- | --- | --- | --- | --- | --- | --- | --- | --- | --- | --- | --- | --- |
| 16SB | - | **-0.79** | 0.35 | **0.95** | 0.06 | -0.24 | **0.73** | **0.74** | 0.05 | 0.35 | -0.77 | 0.32 |
| 16SA | **-0.79** | - | -0.62 | **-0.90** | -0.50 | 0.32 | **-0.86** | **-0.85** | -0.33 | -0.30 | 0.48 | -0.28 |
| 18SF | 0.35 | -0.62 | - | 0.33 | 0.40 | **-0.78** | 0.60 | 0.54 | 0.24 | 0.19 | -0.22 | -0.12 |
| ORR | **0.95** | **-0.90** | 0.33 | - | 0.25 | -0.22 | **0.95** | **0.85** | 0.10 | 0.22 | -0.45 | 0.15 |
| %N removal | 0.06 | -0.50 | 0.40 | 0.25 | - | **-0.95** | 0.19 | 0.15 | **0.85** | -0.18 | -0.46 | -0.15 |
| Redox potential | -0.24 | 0.32 | **-0.78** | -0.22 | **-0.95** | - | -0.34 | -0.36 | **-0.94** | 0.19 | 0.24 | 0.22 |
| Voltage production | **0.73** | **-0.86** | 0.60 | **0.95** | 0.19 | -0.34 | - | **0.90** | 0.10 | 0.25 | -0.54 | 0.21 |
| CE | **0.74** | **-0.85** | 0.54 | **0.85** | 0.15 | -0.36 | **0.90** | - | 0.11 | 0.26 | -0.50 | 0.17 |
| pH | 0.05 | -0.33 | 0.24 | 0.10 | **0.85** | **-0.94** | 0.10 | 0.11 | - | -0.22 | -0.22 | -0.20 |
| Temperature | 0.35 | -0.30 | 0.19 | 0.22 | -0.18 | 0.19 | 0.25 | 0.26 | -0.22 | - | **-0.75** | **0.75** |
| Conductivity | **-0.77** | 0.48 | -0.22 | -0.45 | -0.46 | 0.24 | -0.54 | -0.50 | -0.22 | **-0.85** | - | **-0.75** |
| Suspended solids | 0.32 | -0.28 | -0.12 | 0.15 | -0.15 | 0.22 | 0.21 | 0.17 | -0.20 | **0.75** | **-0.75** | - |

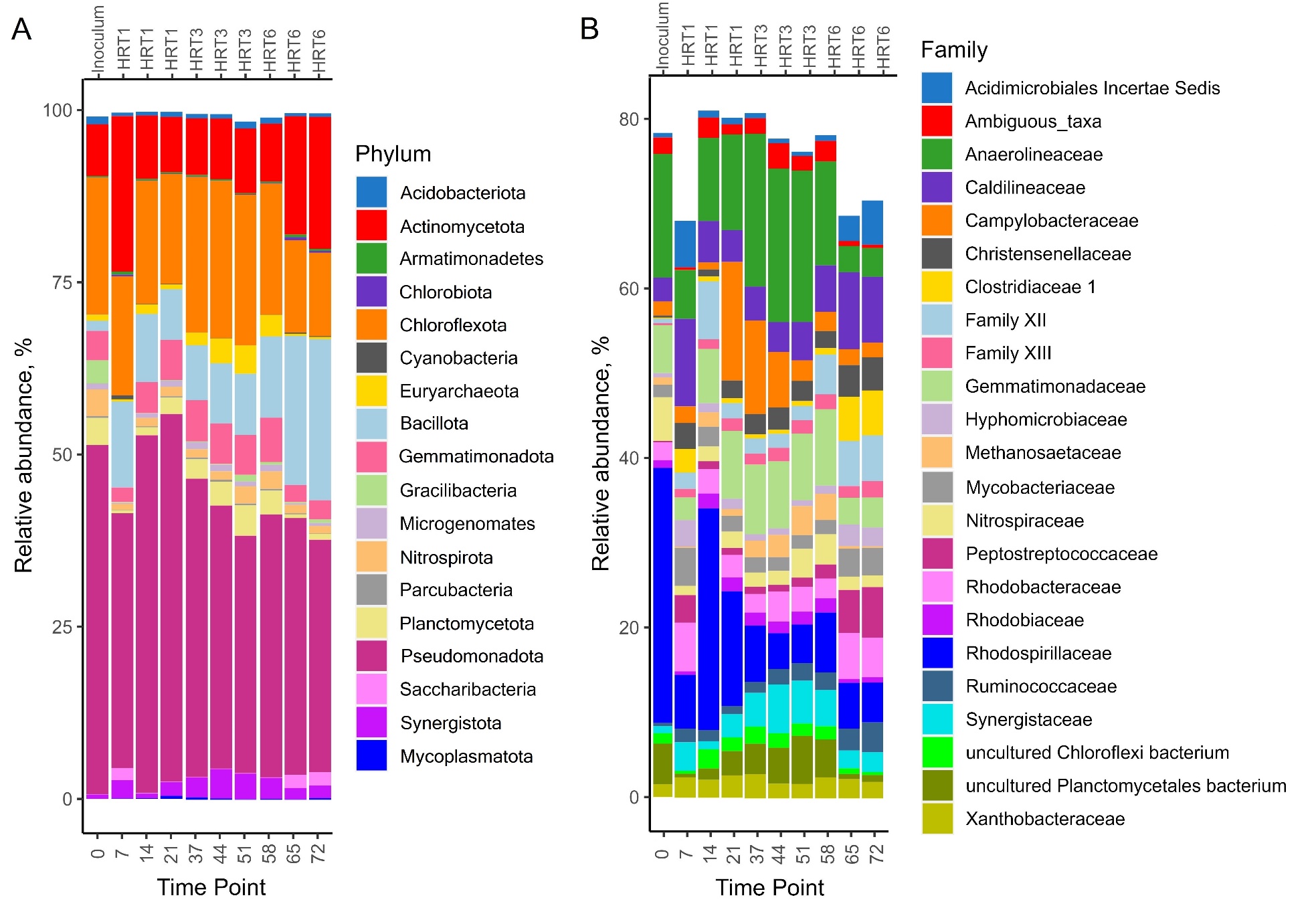

**Supplementary Fig. S3.** Relative abundance of prokaryotic ASVs at the phylum (A) and family (B) taxonomic levels in the original inoculum and at different time points during the experimental period. Three consecutive HRTs were examined: 1 day (HRT1), 3 days (HRT3), and 6 days (HRT6).

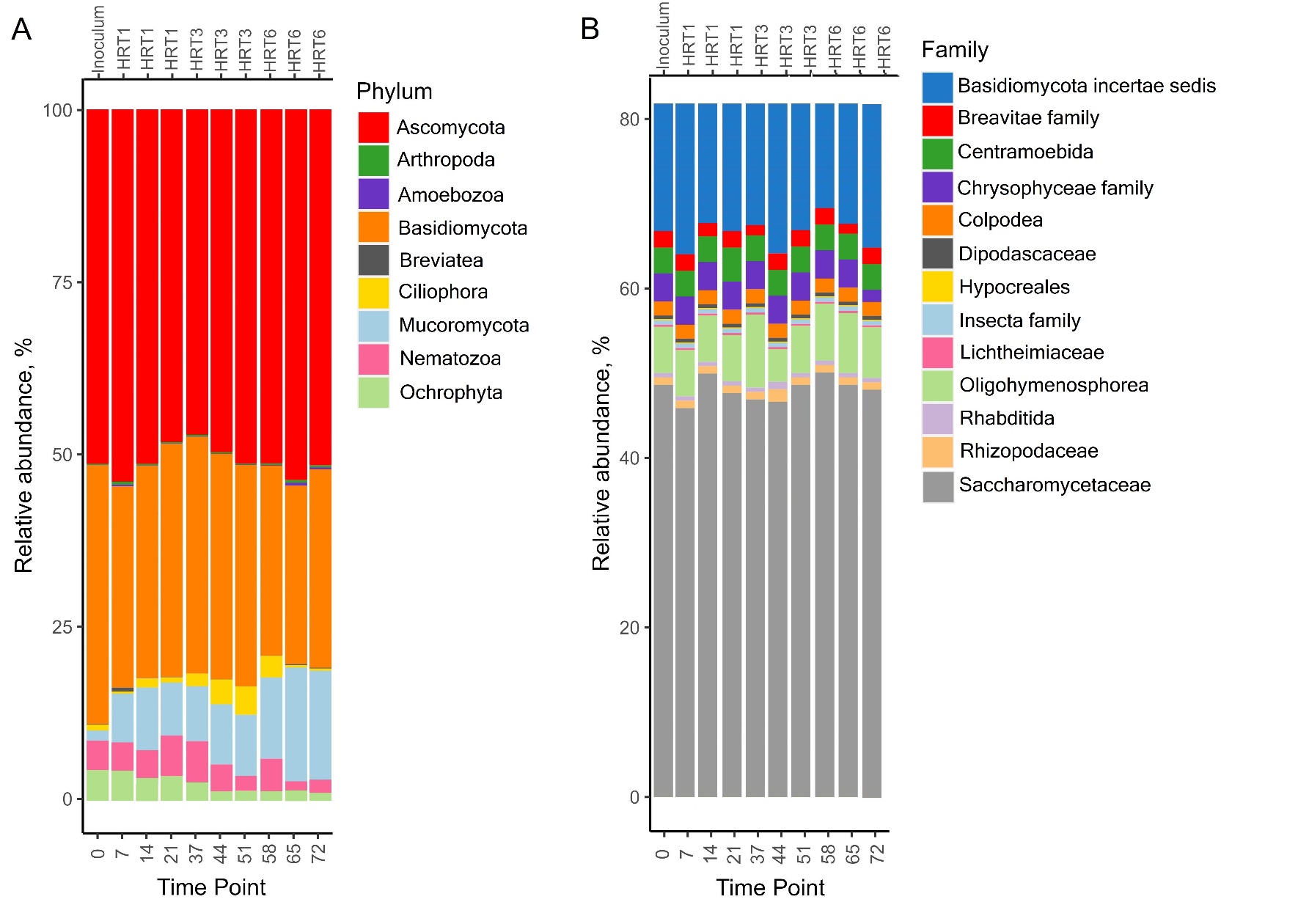

**Supplementary Fig. S4.** Relative abundance of eukaryotic ASVs at the phylum (A) and family (B) taxonomic levels in the original inoculum and at different time points during the experimental period. Three consecutive HRTs were examined: 1 day (HRT1), 3 days (HRT3), and 6 days (HRT6).

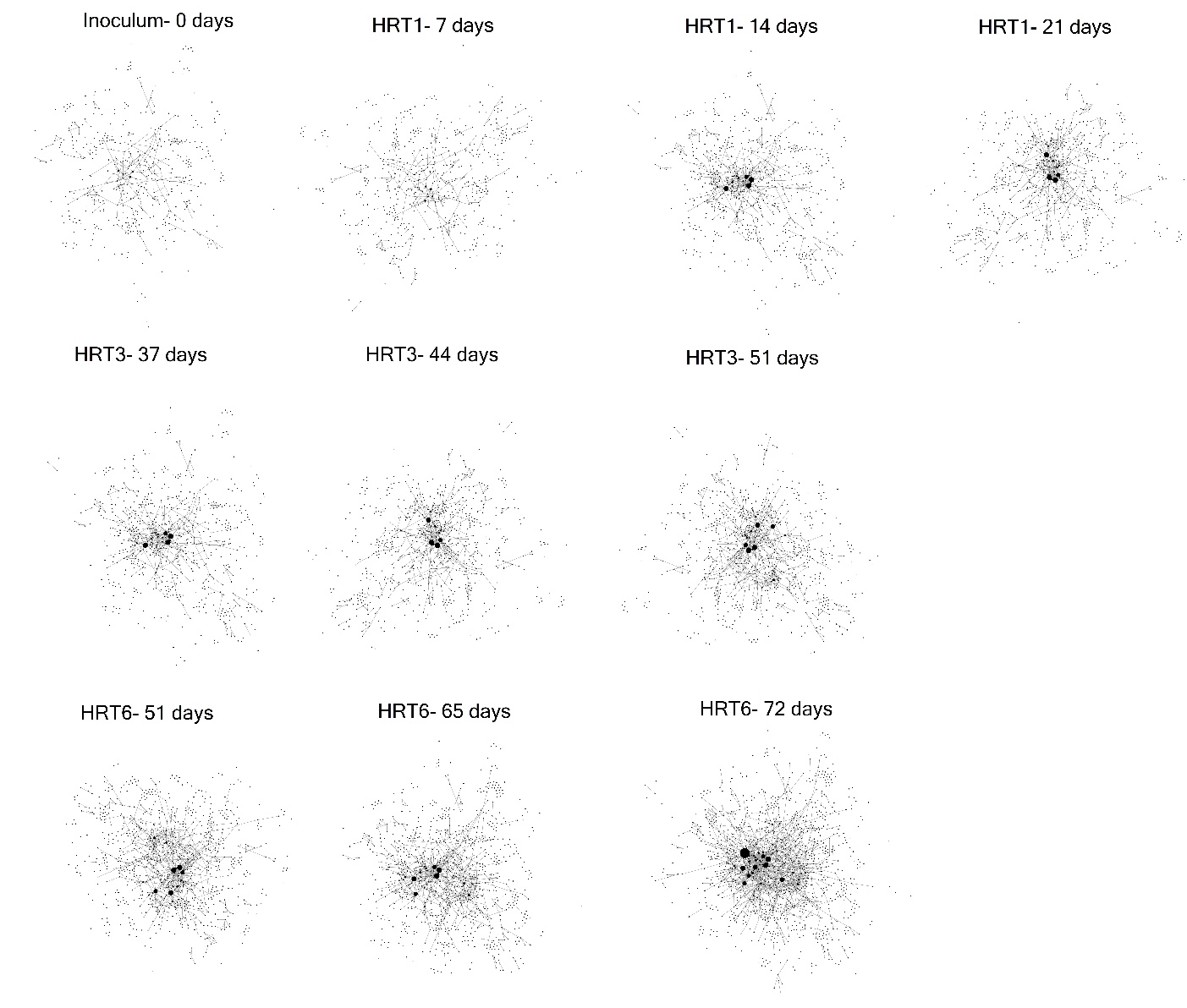

**Supplementary Fig. S5.** Co-occurrence networks of prokaryotic microbial communities in the original inoculum and at different time points during the experimental period. Three consecutive HRTs were examined: 1 day (HRT1), 3 days (HRT3), and 6 days (HRT6). Nodes indicate ASVs and edges represent significant co-occurrence relationships (Spearman’s ρ > 0.75 and *p* ≤ 0.05). Network properties and their ecological relevance are described in more detail in Supplementary Table S3.

**Supplementary Table S4.** Relevant properties of the co-occurrence networks of prokaryotic microbial communities displayed in Supplementary Fig. S5 at different time points. For each row, different letters indicate significant differences between treatments (Tukey's HSD, *p* ≤ 0.05).

|  | | | | **Inoculum** | **HRT1** | | | **HRT3** | | | **HRT6** | | |
| --- | --- | --- | --- | --- | --- | --- | --- | --- | --- | --- | --- | --- | --- |
|  |  |  |  | **Time Points** | | | | | | | | | |
| **Category** | **Metric** | **Definition** | **Ecological relevance** | **0** | **7** | **14** | **21** | **37** | **44** | **51** | **58** | **65** | **72** |
| Size | Nodes | Each node represents a bacterial ASV | Larger networks contain a greater number of interacting (co-occurring or co-excluding) ASVs | 401e | 430d | 481c | 485c | 511b | 520b | 535b | 560a | 568a | 580a |
| Size | Edges | Edges indicate significant co-occurrence or co-exclusion relationships. | Co-occurrence could represent a number of ecological interactions, from predator-prey relationships to commensalism to shared ecological niches (Faust and Raes 2012). Co-exclusion may represent competition or inhibition | 2012e | 2589d | 2876c | 2890c | 3050b | 3078b | 3100b | 3202a | 3210a | 3219a |
| Degree | Mean degree | Degree refers to the number of edges a given node has. Mean degree is the average degree across all nodes in a network (Berry and Widder 2014) | Higher mean degree indicates more co-occurrence or co-exclusion relationships per ASV | 7.2e | 7.8d | 8.5c | 8.9c | 10.3b | 10.6b | 10.9b | 11.4a | 11.7a | 12.1a |
| Cohesion | Density | Density is defined as the ratio of the number of edges in a given network to the number of edges possible for that many nodes | High-density networks contain a large proportion of interacting ASVs | 0.25d | 0.28cd | 0.32c | 0.36c | 0.43bc | 0.45b | 0.50ab | 0.53a | 0.54a | 0.59a |
